# The association of human astrovirus with extracellular vesicles facilitates cell infection and protects the virus from neutralizing antibodies

**DOI:** 10.1101/2021.05.17.444590

**Authors:** Baez-N Carlos, Iván R. Quevedo, Susana López, Carlos F. Arias, Pavel Iša

## Abstract

Viral gastroenteritis has a global distribution and represents a high risk for vulnerable population and children under 5 years because of acute diarrhea, fever and dehydration. Human astroviruses (HAstV) have been identified as the third most important cause of viral gastroenteritis in pediatric and immunocompromised patients. Furthermore, HAstV has been reported in biopsies taken from patients with encephalitis, meningitis and acute respiratory infection, yet it is not clear how the virus reaches these organs. In this work we tested the possibility that the released astrovirus particles could be associated with extracellular vesicles. Comparison between vesicles purified from astrovirus- and mock-infected cells showed that infection with HAstV Yuc8 enhances production of vesicles larger than 150 nm. These vesicles contain CD63 and Alix, two markers of vesicular structures. Some of the extracellular virus was found associated with vesicular membranes, and this association facilitates cell infection in the absence of trypsin activation and protects virions from neutralizing antibodies. Our findings suggest a new pathway for HAstV spread and might represent an explanation for the extraintestinal presence of some astrovirus strains.

**Importance:** Astroviruses are an important cause of diarrhea in children; recently some reports have found these viruses in extra-intestinal organs, including the central nervous system, causing unexpected clinical disease. In this work we found that human astrovirus strain Yuc8 associates with extracellular vesicles, possibly during or after their cell egress. The association with vesicles seems to increase astrovirus infectivity in less susceptible cells, and renders virus particles insensitive to neutralization by antibodies. These data suggest that extracellular vesicles could represent a novel pathway for astrovirus to disseminate outside the gastrointestinal tract.

## Introduction

Astroviruses are considered the third most important cause of viral gastroenteritis in children, as well as in the young of many animal species (1, 2). Moreover, in some mammalian species astroviruses have been associated with different neurological disorders and have been found in biopsies of patients with encephalitis, meningitis or acute respiratory infections (1, 2). Given that mammalian astroviruses are considered intestinal viruses, the central question is: “How could astroviruses get into the central nervous system and respiratory tract?” (3).

From the structural point of view, astroviruses are small non-enveloped viruses, forming the family *Astroviridae*. They contain a single-stranded, positive sense RNA (ssRNA+) genome whose length ranges, in the case of mammalian astroviruses, from 6.1 to 6.8 kb. The astrovirus genome is organized into three open reading frames, named ORF1a, ORF1b and ORF2, which encode non-structural (ORF1a and ORF1b) and structural (ORF2) viral proteins (4, 5). Astrovirus cell entry is not completely understood, and the virus cell surface receptor is unknown, although the fact that the susceptibility of different cell lines to infection with astrovirus depends on the viral serotype (6–8), suggests that there could be more than one receptor. Astrovirus enters into cells by clathrin-mediated endocytosis and it seems that entry process follows a classical route into late endosomes (4, 9). During maturation, the astrovirus particles are subjected to distinct proteolytic processes. First, the capsid protein VP90 of the newly assembled astrovirus particles is cleaved intracellularly by caspases to give immature virions composed by the viral protein VP70. This cleavage is associated with the release of the viral particles from the infected cell (10, 11). Then, once in the extracellular medium, the virion is processed by trypsin-like extracellular proteases to render infectious, mature virions, composed by the final protein products VP27 and VP34 (12, 13).

One of the less characterized phases of the astrovirus replication cycle, is cell egress. It has been proposed that astrovirus release is a non-lytic process, during which the extracellular virions appear to be associated with membranous structures (11, 14). In this regard, it is of interest that the cell exit of different viruses has been associated with extracellular vesicles (EV) (15–17). EV are a heterogeneous group of small vesicles with a lipidic bilayer, ranging from 50 nm to 1,000 nm of diameter (18). These vesicles are secreted by different types of cells and can be isolated from conditioned media of cultured cells, as well as from virtually any type of body fluid, including blood, urine, ascites, bronchoalveolar lavage, saliva and cerebrospinal fluid (19, 20). There are different types of EV, with exosomes being the better characterized, having a diameter of around 50 to 150 nm, and also well studied microvesicles with diameter around 50 to 1,000 nm. Exosomes originate from the endosomal compartment by fusion of multivesicular bodies with the plasma membrane, while microvesicles originate from the plasma membrane by outward budding (21).

Viral infections affect cell physiology, as well as many cellular processes, including protein synthesis and degradation (22, 23), intracellular trafficking and vesicle secretion (16, 24, 25). In the last few years the evidence regarding the interaction between EV and different types of viruses (26–28) has accumulated. Particularly, several positive-sense ssRNA viruses, like hepatitis C virus (HCV) and hepatitis E virus (HEV), have been found to associate with EV or to use the mechanism of EV biogenesis as an egress pathway (29–32). In addition, DNA viruses like HSV-1 (33) and JC polyomavirus (34) also have been observed interacting or being released with EV.

Given the possibility that EV could be involved in the human astrovirus (HAstV) cell egress, we tested the possibility that astrovirus particles could be released in association with this type of vesicles. To characterize the possible interaction between EV and the virus, Caco-2 cells were infected with the Yuc8 strain of HAstV and EV were purified from the cell culture media by differential centrifugation coupled to polyethylene glycol 6000 (PEG) precipitation and affinity magnetic sorting. Our results suggest that astrovirus infection stimulates the secretion of EV and astrovirus particles seem to associate with EV. These vesicle-associated viruses acquire the ability to infect cells in the absence of trypsin activation. Also, viral particles associated with EV were refractory to the effect of neutralizing antibodies, suggesting that EV are able to protect the virions from this interaction.

## Materials and methods

### Cell lines, virus, reagents and antibodies

Human colon adenocarcinoma cells (Caco-2), and rhesus monkey epithelial cells (MA104), were obtained from American type culture collection (ATCC, Manassas, VA, USA). Dulbecco modified Eagle medium - high glucose (DMEM) was purchased from Sigma Aldrich (San Luis, MI, USA), while Advanced-DMEM (A-DMEM), fetal bovine serum (FBS) and trypsin were from Gibco (Thermo Fisher Scientific, USA). Triton X-100 was acquired from Boehringer Mannheim, (Germany), whereas Polyethylene glycol 6000, soybean trypsin inhibitor and Minimum Essential Medium (MEM) were acquired from, Sigma-Aldrich (San Luis, MI, USA). Formaldehyde was obtained from J.T. Baker, (USA), and MagCapture™ exosome isolation kit PS was from FUJIFILM Wako Pure Chemical Corporation (Osaka, Japan). Human astrovirus serotype 8, strain Yuc8 was originally isolated in our laboratory (35). Polyclonal rabbit antibody specific for Yuc8 virus (anti-Yuc8) was prepared in our laboratory (11). Rabbit polyclonal antibodies specific for anti-CD63 and anti-Alix were acquired from Santa Cruz (Santa Cruz Biotechnology, CA, USA), and Aviva Systems Biology (Aviva Systems Biology, CA, USA) respectively, while monoclonal antibody specific to anti-protein disulfide isomerase (PDI, clone 1D3) was obtained from Enzo Life Sciences, Inc (C. Mexico, Mexico). Anti-rabbit peroxidase conjugated antibody was from KPL (MD USA), and protein A, peroxidase conjugate was from Sigma Aldrich (Sigma Aldrich).

### Cell culture and viral propagation

Caco-2 cells were cultured in DMEM supplemented with non-essential amino acids and 15% heat-inactivated FBS, in a 10% CO_2_ atmosphere at 37°C (14). MA104 cells were grown in A-DMEM, supplemented with 5% FBS, at 37°C in a 5% CO_2_ atmosphere (36).

A working stock of human astrovirus serotype 8, strain Yuc8 (35), was prepared as previously described (37). The virus was activated just prior to infection with 200 μg/mL of trypsin, for 1 hour at 37 °C, followed by inactivation with 200 μg/mL of soybean trypsin inhibitor. Before infection, Caco-2 cell monolayers were washed with MEM and incubated with activated virus for 1 hour at 37 °C. Then, cell monolayers were washed twice with MEM, to remove non-adsorbed virus. Finally, MEM was added to the cells, and infection was left to proceed for 48 hours at 37 °C.

Astrovirus particles were purified essentially as described previously (38). Briefly, Caco-2 cells were infected with HAstV serotype 8 (Yuc8) at an MOI of 5 as described above and the infection was left to proceed for 48 hours. After this time cells were detached, and frozen and thawed three times. Then, cellular lysate was clarified by centrifugation at 2,000 g for 10 minutes and then passed through a 0.45 μm filter (Milipore). Filtered supernatants were pelleted at 60,000 g for 16 hours at 4 °C in a SW28 Ti rotor (Beckman), and the resulting pellet was resuspended in TNE buffer (50 mM Tris-HCl [pH 7.4], 0.1 M NaCl, 10 mM EDTA). This suspension was adjusted to 0.5% v/v with octyl glucoside in TNE buffer and incubated for 30 minutes on ice. Finally, virus was pelleted through a 30% w/v sucrose cushion in TNE buffer for two hours at 200,000 g in a SW55 Ti Beckman rotor. The pelleted viral particles were resuspended in TNE buffer.

### Viral infectivity assay

Viral titers were determined by immuno-peroxidase staining to detect infectious focus forming units (FFU) as described previously (11, 39). In brief, Caco-2 cells were cultured to confluence in 96 wells plates and washed with serum-free MEM before infection. Viral samples were activated with trypsin (200 μg/mL), for 1 hour at 37 °C, soybean trypsin inhibitor was added (200 μg/mL), and serial fold dilutions of activated viral samples were performed. Diluted samples were added to each well and let to adsorb for 1 hour at 37 °C. After adsorption period, the virus inoculum in each well was removed, cells were washed twice with MEM and infection was left to proceed in fresh MEM for 18 hours at 37°C. Cells were fixed for 20 minutes with 2% formaldehyde in phosphate-buffered saline (PBS), then they were washed three times with PBS and permeabilized by a 15 minutes incubation with 0.2% Triton-X100 solution in PBS. Finally, cells were washed again three times with PBS and incubated with a polyclonal rabbit anti-Yuc8 overnight at 4°C. Next day cells were washed out three times with PBS and incubated with peroxidase conjugated protein A for 2 hours at 37°C. After washing protein A, infected cells were revealed by carbazole precipitation and FFU were counted.

### Kinetics of viral release

Caco-2 cells were grown to confluence in 24 wells plates. Cell monolayers were washed twice with MEM and infected with activated HAstV Yuc8 strain (at an MOI of 5). Supernatants were harvested at three hours intervals starting at 12 hours post infection (hpi) until 24 hpi, and centrifuged for 5 minutes at 500 g to separate cellular debris. At the same time, MEM was added to cellular monolayers and cells were lysed by two cycles of freeze-thaw. Infectious viral particles associated to cells and present in supernatants were determined by an immune-peroxidase assay as described above. Before trypsin activation, samples were incubated for 30 minutes at 37°C with MEM or with 0.1% Triton X-100 diluted in MEM.

### Purification of extracellular vesicles

Caco-2 cells, grown to confluence in 150 cm^2^ flasks, were washed twice with MEM and infected with trypsin activated HAstV Yuc8 at an MOI of 5. As a control, cells were mock infected using an identical protocol without virus. Supernatants were harvested at 18 hpi and processed by differential centrifugation essentially as described before (40, 41). Briefly, supernatants were centrifuged at 500 g for five minutes to obtain pellet 1 (P1), and the supernatant was again centrifuged at 2,000 g for 30 minutes, obtaining the pellet 2 (P2). The remaining supernatant was centrifuged at 20,000 g for one hour, producing pellet 3. Finally, the last supernatant was mixed with an equal volume of a solution of 16% polyethylene glycol 6000 (PEG), 1 M sodium chloride and left overnight at 4°C. The mixture was then centrifugated at 10,000 g for one hour, yielding pellet 4. As proposed by a theoretical analysis of sedimentation (42), the purified fraction in pellet 3 was considered to contain large extracellular vesicles (LEV), while the fraction of pellet 4 contains small extracellular vesicles (SEV). All centrifugations were performed at 4 °C and all pellets were resuspended in sterile PBS. Virus titer in purified fractions was determined by immune-peroxidase assay, with and without TX-100 treatment as described above for supernatants in the assays of viral release kinetics.

For some experiments, in order to remove possible contaminants (i.e., free contaminating virions or protein aggregates) from purified vesicles in LEV or SEV fractions (pellets 3 and 4, respectively), the vesicle fractions were additionally purified using the MagCapture™ exosome isolation kit PS, according to the manufacturer protocol.

### Immunodetection of cellular and viral proteins

The fractions purified by differential centrifugation from supernatants of infected and mock-infected Caco-2 cells were mixed with Laemmli sample buffer (50 mM Tris, pH 7.5, 2% SDS, 2% β-mercaptoethanol, 10 mM EDTA and 0.1% bromophenol blue), boiled for 5 min and the proteins were separated by sodium dodecyl sulfate-polyacrylamide gel electrophoresis (SDS-PAGE). Proteins were transferred to a nitrocellulose membrane (Millipore, Bedford, MA, USA). Membranes were blocked with 5% non-fat dried milk in PBS. The proteins of interest were detected with specific primary antibodies followed by incubation with secondary peroxidase-conjugated reagents. Primary antibodies were incubated with membranes overnight at 4°C, washed three times with PBS 0.1% Tween (PBS-T) and incubation continued with peroxidase conjugated secondary antibody or protein A for 90 min, at room temperature. After these incubations the membranes were washed again with PBS-T and proteins were visualized by Western Lightning Chemiluminescence Reagent Plus (Perkin Elmer).

### Infectivity associated to the extracellular vesicles

Fractions were purified from supernatant of astrovirus infected Caco-2 cells by differential centrifugation coupled with isolation with magnetic beads (MagCapture™ exosome isolation kit PS), as described above. Vesicle containing fractions were diluted in MEM and added to Caco-2 and MA104 cells, grown to confluence in 96 wells plates, washed twice with MEM before addition. The fractions were subjected to the following treatments before adsorption: with 0.1% Triton X100 for 30 minutes at 37°C; or were preincubated with an anti-Yuc8 neutralizing antibody for 1 hour at 37°C; or by an incubation with 0.1% Triton X-100 for 30 minutes at 37°C, followed by neutralization with anti-Yuc8 antibody (1 hour at 37°C). Control, non-treated samples were incubated in the same conditions but using an equivalent volume of FBS-free MEM instead of Triton X-100 and/or anti-Yuc8 antibody. Vesicles were left to adsorb to cells during 2 hours at 37°C, then cells were washed, fresh medium was added and the infection was left to proceed for 18 hours.

To test the capacity of purified EV to promote infection with externally bound virus particles, LEV and SEV vesicles were purified from non-infected Caco-2 cells as described above, including the isolation step with the MagCapture™ exosome isolation kit PS. These vesicles were then incubated for 1 hour at 37 °C with a known amount of non-activated purified HAstV Yuc8. After incubation, the vesicles were treated as described above (0.1% Triton X-100 for 30 minutes at 37°C, anti-Yuc8 neutralizing antibody for 1 hour at 37°C or 0.1 % Triton X-100 followed by neutralization with anti-Yuc8 antibody). After the treatments, vesicles with virus were added to Caco-2 or MA104 cells grown in 96 wells, and incubated during 2 hours at 37°C. After this time cells were washed, and infection was left to proceed for 18 hours. As control, an equal amount of the identical purified trypsin activated or nonactivated astrovirus was used in the same conditions without EV incubation. Infected cells were counted in selected area, of two wells per sample using 20X lens. Images were acquired with 10X lens in a Nikon Diaphot 300 microscope.

### Transmission Electron Microscopy

LEV and SEV vesicular fractions were purified from infected Caco-2 cells as described above, using differential centrifugation coupled to isolation with the MagCapture™ exosome isolation kit PS. Purified fractions and non-activated purified virus particles were bound on carbon vaporized copper grids covered with formvar and negatively stained with 3% uranyl acetate. Images were acquired using a Zeiss Libra 120 electron microscope operating at 80 KV coupled with a GATAN Multiscan 600HP 794 CCD camera.

### Nanoparticle Tracking Analysis

Nanoparticle tracking analysis (NTA), was conducted using a NanoSight NS300 (Malvern Instruments Ltd., Worcestershire, UK) to assess the hydrodynamic diameter of non-activated virus particles and vesicles purified by differential centrifugation from infected and non-infected cells supernatants. Purified fractions were analyzed after diluting the samples in sterile and microfiltered PBS (1:100-200 in the case of vesicles and 1:1000 in case of purified virus particles). For each condition 5 videos of one-minute length each, were recorded sizing 20-40 particles/frame and analyzed using the NanoSight NTA 3.1 software (43). This technique uses dynamic light scattering to measure the diffusion coefficient of particles moving under Brownian motion and converts it to hydrodynamic diameter using the Stokes-Einstein equation (44). A blank of sterile filtered PBS was used for particle calculations in every measurement, and after each measurement the flushing lines were thoroughly washed three times to prevent contamination.

### Statistical Analysis

Statistical analysis of the obtained results was performed using the GraphPad prism 5.0 software (GraphPad Software, Inc.), with an interval of confidence of 95%.

## Results

### HAstV Yuc8 titer in detergent-treated media increases with time of infection

It has been previously observed that a fraction of the astrovirus particles produced in infected cells floats to low-density fractions when separated by density centrifugation, suggesting that they interact with membranous structures (14). To characterize the possible association of astrovirus particles with membranes in the cell culture media, we evaluated the kinetics of astrovirus release in Caco-2 cells. Media from astrovirus infected cells were collected at different time points after infection (from 12 to 24 hpi), and the virus titer was determined. The infectivity of the virus in the supernatant was activated with trypsin after the samples were treated or not with detergent (Triton X-100). Under these conditions, if astrovirus particles were associated with vesicles or membranes, the detergent treatment would release the virions, leading to an increase in viral titer as compared to the titer of samples not treated with detergent. We observed the virus present in the supernatant starting at the first time point analyzed (12 hpi), without any significant change in the titer after Triton X-100 treatment (p>0.05) (Fig. 1), similar to previously published results (10). The titer of virus present in the media not treated with detergent showed little increase from 15 to 24 h post infection, however, the viral titer increased considerably in the supernatant after detergent treatment, reaching almost twice as much infectivity compared to non-treated samples, at later time points (Fig. 1). No significant cellular damage was detected at the different time points (being under 10% of total LDH in both mock and Yuc8 infected cells), as determined by an LDH assay (results not shown). These results suggest that astrovirus particles are released from infected cells before appreciable cell lysis, and that they could be associated with detergent soluble structures in the extracellular medium. In all the subsequent experiments shown here, media were harvested at 18 hpi.

**Figure 1.**
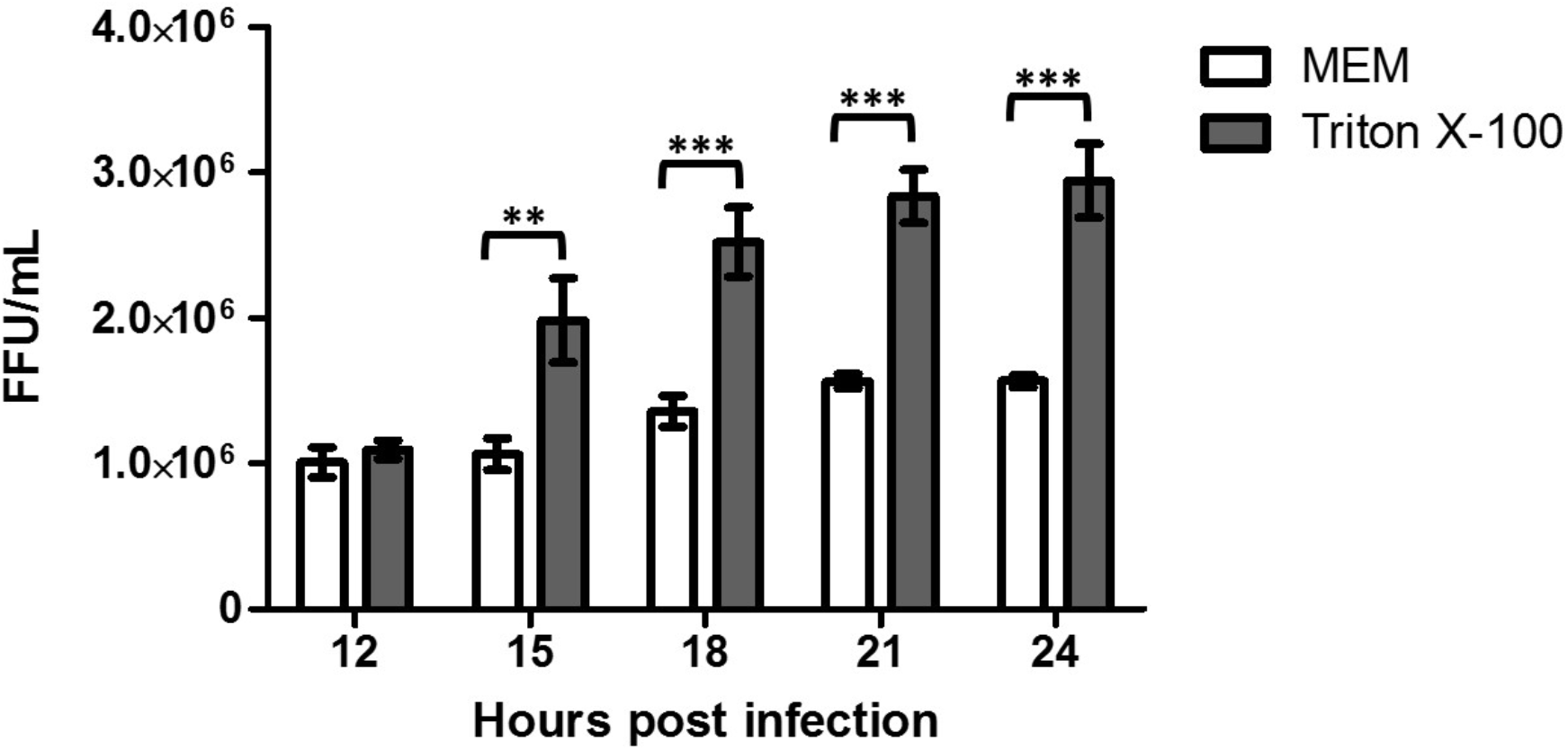
Kinetics of astrovirus release from infected Caco-2 cells. Caco-2 cells were infected with astrovirus Yuc8 at MOI of 5 and supernatants were collected at different time points post infection (from 12 to 24 hours). Viral titer was determined with (grey bars) and without (white bars) 0.1% Triton X-100 treatment before trypsin activation. The results represent the mean focus forming units (FFU) per milliliter (mL) ± standard error of the mean of three independent experiments done in duplicate. Statistical analysis was done with two-way ANOVA, p value *<0.05; ** p<0.01, *** p<0.001.

### Astrovirus infection increases the secretion of extracellular vesicles from Caco-2 cells

To characterize the effect of astrovirus infection on the production of EV in Caco-2 cells, supernatants from infected or mock-infected cells cultured in serum-free MEM were harvested at 18 hpi and processed by differential centrifugation. Initially, detached cells were pelleted at 500 g for 5 min, getting pellet 1 (P1). Pellet 2 (P2) was obtained by centrifugation of the remaining supernatant at 2,000 g for 30 min. We expected this fraction to contain very large vesicles and some cell debris and organelles. Pellet 3 was obtained by centrifugation of the remaining supernatant at 20,000 g for 1 h, to collect large extracellular vesicles (LEV), theoretically calculated to be over 122 nm (42). Finally, pellet 4 was obtained by overnight precipitation of remaining vesicles and particles in the remaining supernatant by 8% PEG 6000 and 0.5 M NaCl, followed by centrifugation at 10,000 g for 1 h, producing small extracellular vesicles fraction (SEV), calculated to be under 170 nm (41, 42). As consequence, we expect some size overlapping between LEV and SEV fractions.

An equal portion of each pellet fraction was analyzed after SDS-PAGE. By silver staining of the gel, it was clear that the amount of total proteins present in each fraction was increased in astrovirus-infected cells (Fig. 2A). The presence of different cellular markers in the pelleted fractions was analyzed by immunoblotting; EV specific markers tested were CD63 and ALIX, while endoplasmic reticulum associated PDI protein was used as non-EV associated protein control. In fractions P1 and P2, which probably contain cells, cell debris and large vesicles, all proteins markers were observed, and their presence also has increased after astrovirus infection. Interestingly, the LEV and SEV fractions purified from astrovirus-infected Caco-2 cells showed a higher content of EV specific proteins (ALIX and CD63) as compared to mock-infected cells (Fig. 2B), presumably representing larger amounts of EVs. (Fig. 2B).

**Figure 2.**
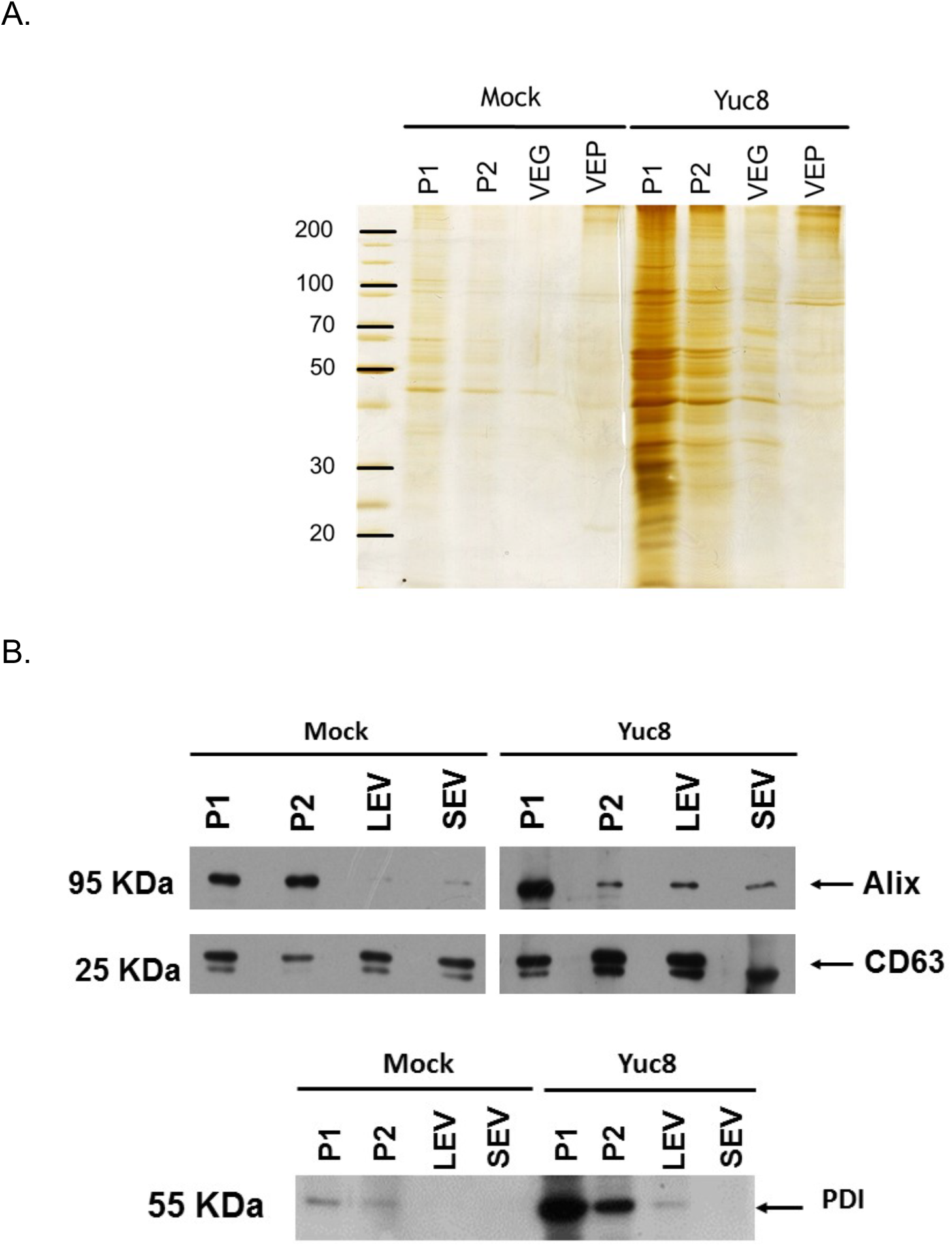
Identification of vesicular markers in vesicles purified from Caco-2 cells. Caco-2 cells were infected or mock-infected with astrovirus strain Yuc8 (MOI 5) and 18 hours post infection the supernatant was collected. Different fractions were purified by differential centrifugation; pellet 1 was obtained by centrifugation at 500 g (P1), pellet 2 obtained by centrifugation at 2,000 g (P2), fraction containing large extracellular vesicles (LEV) was obtained after 20,000 g centrifugation and final small extracellular vesicles (SEV) fraction was obtained by precipitation with 8% polyethylene glycol 6000, 0.5M NaCl. (A) The same volume of each pelleted fraction was separated by SDS-PAGE and proteins were detected by silver staining. (B) Samples were resolved on SDS-PAGE and analyzed by western blotting, using antibodies specific for CD63 and Alix as vesicle markers and protein disulfide isomerase (PDI) as endoplasmic reticulum protein to assess preparation contamination. Immunoblots are representative of five independent experiments.

To quantitate the concentration and size of the purified vesicles more precisely, the pellet 3 (LEV fraction) and the fraction purified after PEG 6000 precipitation (SEV fraction) were analyzed by nanoparticle tracking analysis (Fig. 3). There was a clear and significative increase in the vesicle number in the LEV fraction from Yuc8-infected cells (p<0.05), compared to that of mock-infected cells (Fig. 3A and 3C). In the case of the SEV fraction obtained by PEG 6000 precipitation, there was only a small, not significant increase (p>0.05) on the number of vesicles present in preparations obtained from either infected cells or mock-infected cells (Fig. 3B and 3C). These results suggest that astrovirus infection might stimulate the production of EV, particularly those present in the LEV fraction.

**Figure 3.**
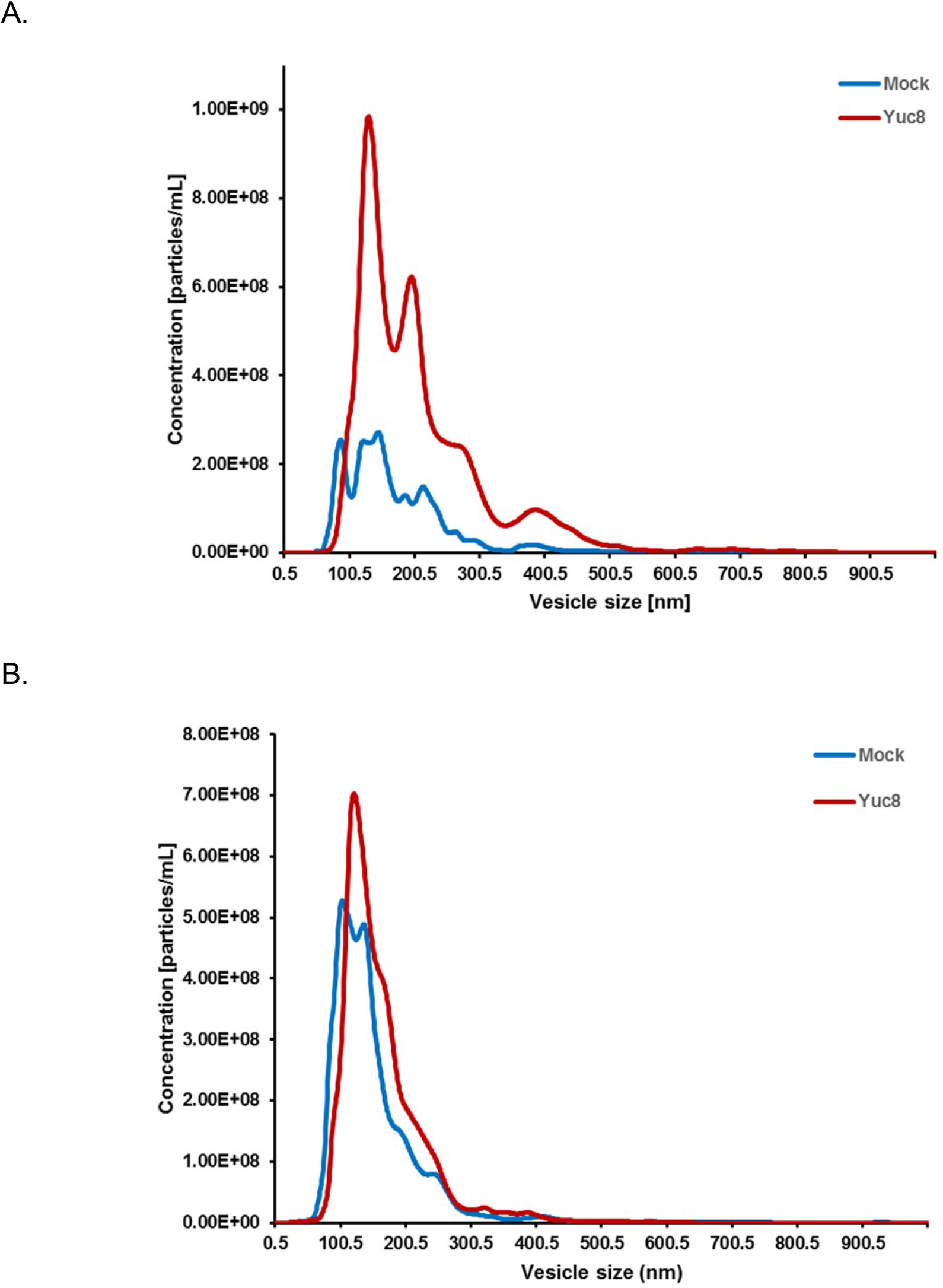
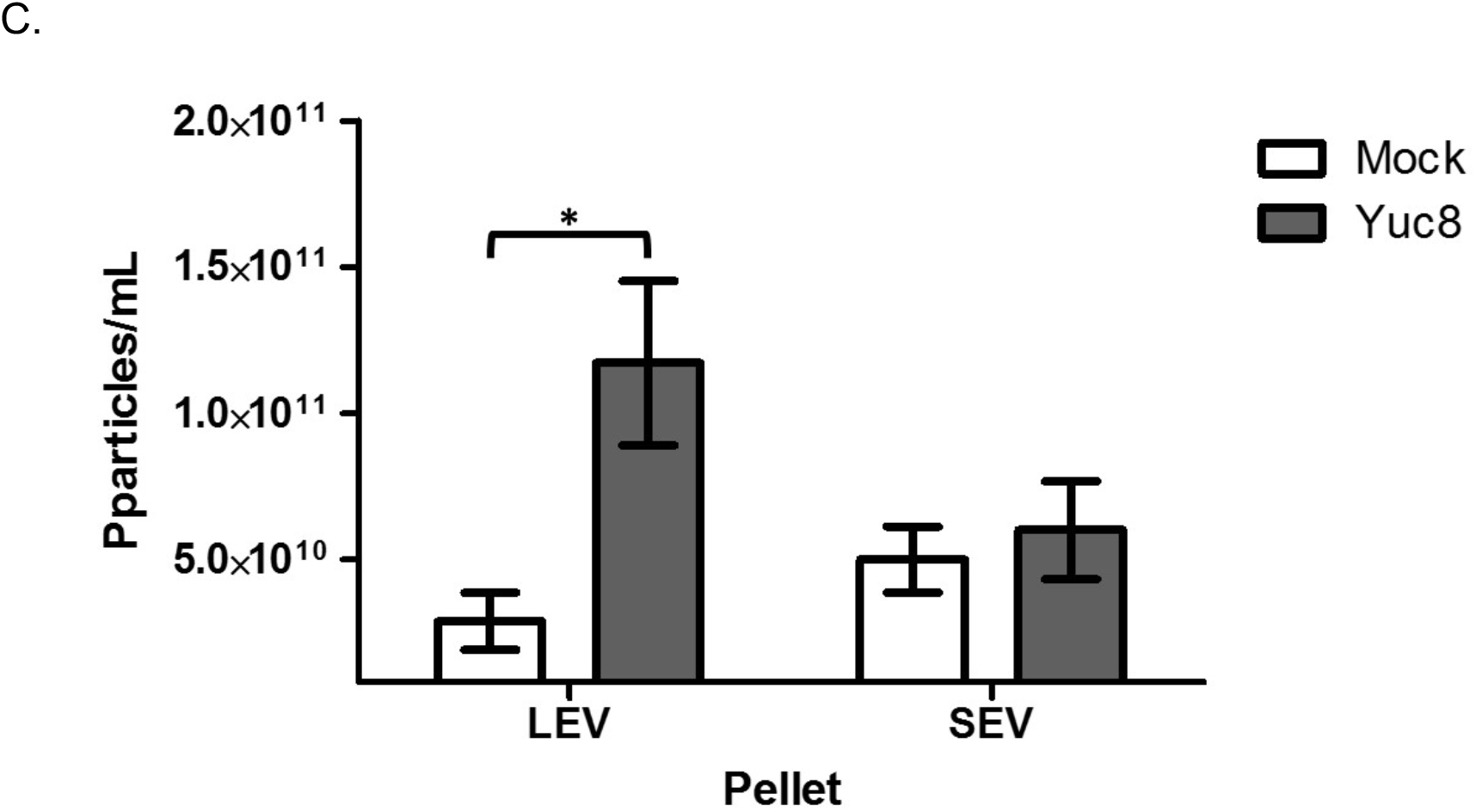
Astrovirus Yuc8 infection stimulates secretion of extracellular vesicles in Caco-2 cells. Confluently grown Caco-2 cells were infected with astrovirus Yuc8 (MOI 5) or mock infected. Supernatants were harvested 18 hours post infection and processed by differential centrifugation. Fractions obtained after pelleting at 20,000 g, corresponding to large extracellular vesicles (LEV) (A) and obtained after PEG 6000 precipitation, small extracellular vesicles (SEV) (B), were resuspended in PBS and used for nanoparticle tracking analysis in the NanoSight NS300. In each experiment five videos were recorded and used for analysis. Distribution of particle-vesicle size (hydrodynamic diameter in nm) and concentration (particles/mL) from 3 to 5 independent experiments are shown. Vesicles purified from mock infected cells are represented by blue line, Yuc8 purified vesicles are represented by red line. (C) Comparison of the mean number of particles present in LEV and SEV fractions shown in A and B. All results are expressed as the mean of the whole concentration of particles ± standard error of the mean of three independent experiments. Statistical analysis was done using two-way ANOVA * p<0.05.

### Astrovirus particles seem to associate with extracellular vesicles

Given the presence of vesicles with different sizes in the cell culture medium, we analyzed whether astrovirus particles were associated with a particular fraction and if an increased infectivity could be observed after treating the different fractions with Triton X-100 before activation of the virus with trypsin. We found that different amounts of infectious viral particles were present in fractions P2, LEV, and SEV; treatment with Triton X-100 before trypsin activation significantly increased virus titer in fractions P2 (p<0.05) and LEV (p<0.01), but not in fraction SEV (Fig. 4A). These observations suggest that a portion of the astrovirus particles could be present inside vesicles or, alternatively, that groups of viral particles could be associated with EV from the outside, and consequently membrane solubilization releases individual particles, increasing virus titer.

**Figure 4.**
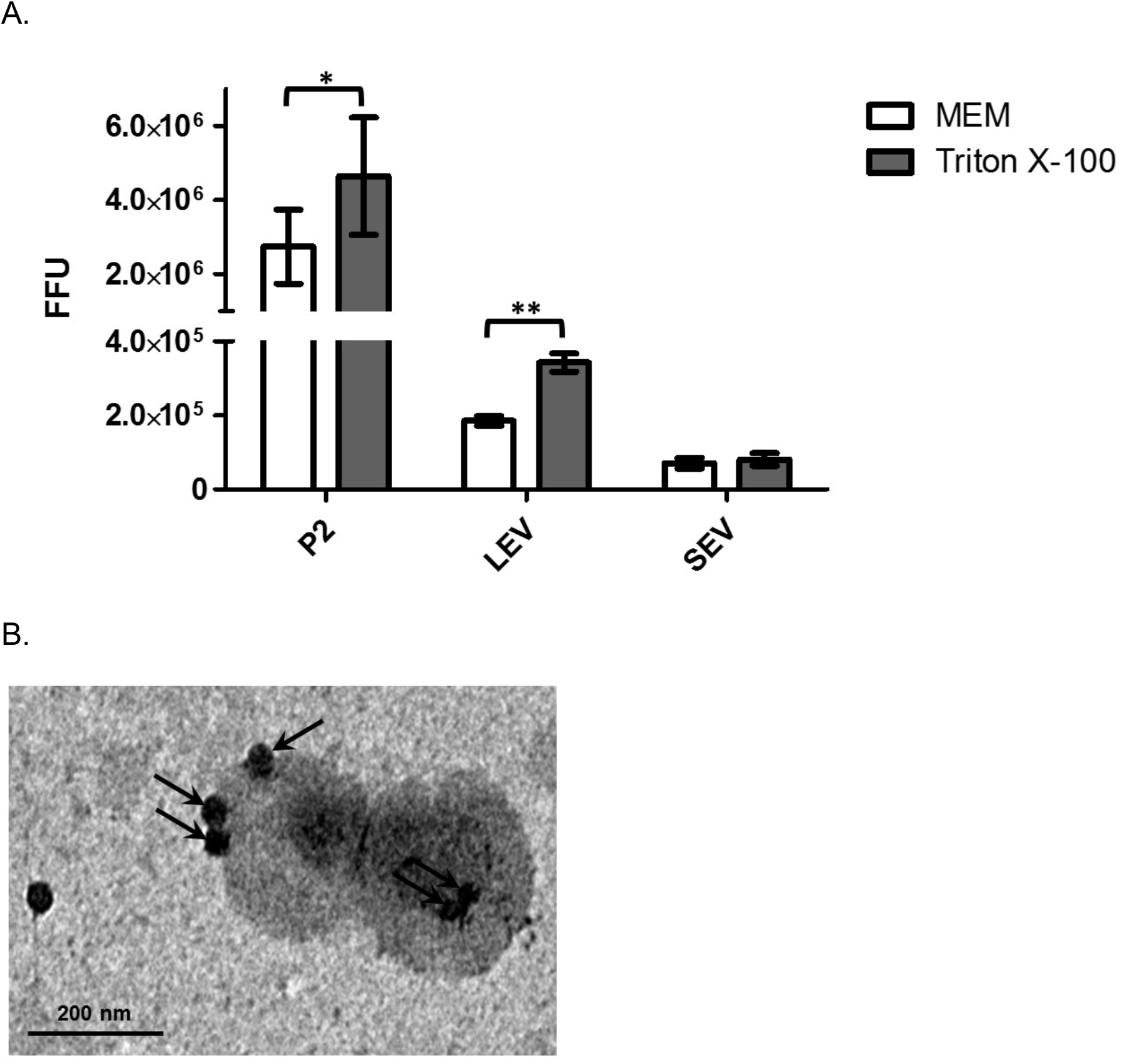
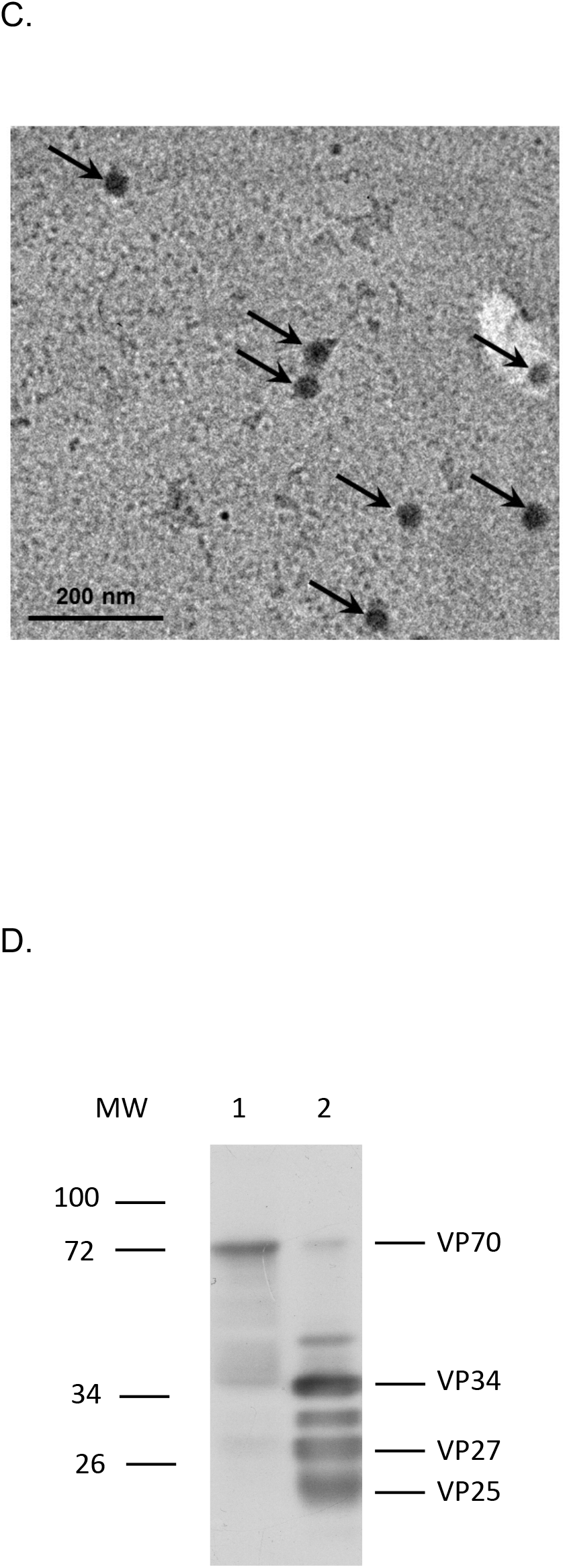
Astrovirus particles associate with large extracellular vesicles. Confluently grown Caco-2 cells were infected with astrovirus Yuc8 (MOI 5) or mock infected. Supernatants were harvested 18 hours post infection and processed by differential centrifugation. Fraction pellet 2 was obtained by centrifugation at 2,000 g (P2), fraction containing large extracellular vesicles (LEV) was obtained after centrifugation at 20,000 g and final small extracellular vesicles (SEV) fraction was obtained by precipitation with PEG 6000, and NaCl after centrifugation at 10,000 g. All fractions were resuspended in same volume (100 μL) of sterile PBS. (A) Purified fractions were trypsin activated, with or without previous incubation with detergent, and titered in Caco-2 cells. Viral content was expressed as total focus forming units. Grey bars represent samples treated with 0.1% Triton X-100 prior trypsin activation; white bars (MEM), represent samples activated with trypsin without Triton X-100 treatment. The mean of viral particles in each sample ± standard error of the mean of three independent experiments are shown. (B) Large extracellular vesicles purified from Yuc8 infected Caco-2 cells as described in A were further clarified by additional isolation using MagCapture™ Exosome Isolation kit PS. One drop of sample was fixed onto carbon vaporized coper grids and negative stained with uranyl acetate. Samples were observed in EFTEM ZEISS Libra 120 electron microscope. Electrodense particles of 30 nm, possibly viral particles, are pointed by black arrows. (C) Purified Yuc8 virions were bound to carbon vaporized coper grids and stained as described in B. Electrodense particles, which resemble astrovirus particles are pointed by arrows. Size bars are shown. (D) Immunoblotting of LEV fraction. Sample used in A, purified from Caco-2 infected cells as described in A, was not-treated (lane 1) or treated (lane 2) with trypsin, and separated in SDS-PAGE, transferred to nitrocellulose membrane and viral proteins were detected using anti-astrovirus polyclonal antibody. Viral proteins are pointed on right hand side, while molecular weight in kilodaltons is shown on left hand side. Images are representative of three independent experiments yielding similar results. Statistical analysis was done with two-way ANOVA, p value *<0.05; ** p<0.01, *** p<0.001.

To determine if there is a direct association between virions and vesicles in the LEV fraction, we analyzed by transmission electron microscopy (TEM) this fraction purified from astrovirus infected CaCo2 cells. The LEV fraction was chosen since the largest increase in virus infectivity when the trypsin activation was done after the Triton X-100 treatment was observed in this fraction. By TEM we found virus-like particles, associated with what appeared to be vesicles (Fig. 4B, pointed by arrows). The electro-dense virus-like particles observed in this micrograph, are similar in form and size (30 nm) to purified astrovirus particles (Fig. 4C), suggesting that they represent bona-fide virus particles associated with membranes. Such viruslike particles were not observed in vesicles present in LEV fraction purified from mock-infected cells (data not shown). Since the infectivity of astroviruses requires activation by proteolytic processing of the VP70 protein precursor, we analyzed by western blot the virus protein composition of the LEV-associated virions. We observed that the virus particles are mainly composed by the VP70 protein (70 KDa) with no evidence of neither VP90 precursor protein, nor any activated viral proteins of 34, 27 or 25 KDa proteins (Fig. 4D).

### Vesicle-associated astrovirus particles are infectious without proteolytic treatment and are protected from antibody neutralization

EVs have an intrinsic capacity to fuse with other cells, and thus to transfer proteins, genetic material, and even viral particles to other recipient cells (16, 21, 24, 45). Using this mechanism, different types of viruses are able to infect otherwise refractory cells. Such is the case of human immunodeficiency virus 1 (HIV-1) (46, 47) and herpes simplex virus 1 (HSV-1) (33). The association with vesicles has also been shown to confer some viruses with resistance to neutralization with specific antibodies [hepatitis A virus (HAV), or HSV-1] (33, 48). To test whether vesicle-associated astrovirus strain Yuc8 is able to infect other cell lines, vesicles present in the LEV and SEV fractions purified from infected Caco-2 cells were added to Caco-2 and MA104 cells. Caco-2 cells were used as fully permissive cell line, while MA104 cells are at least 100 times less permissive to astrovirus Yuc8 infection (8, 49). Before adding to the cell monolayers, the samples were either incubated with 0.1% Triton X-100 for 1 h to disrupt possible membranes; or incubated with polyclonal neutralizing polyclonal antibodies to Yuc8 to neutralize the infectivity of accessible viral particles; or incubated with detergent followed by neutralization with the neutralizing polyclonal antibodies in order to neutralize all viral particles present. As control, fractions were only incubated in MEM.

The results of these assays show that the LEV and SEV vesicle-associated astrovirus viral particles were able to infect both Caco-2 and MA104 cells (Fig. 5A and B), while viral infection was completely abolished after membrane solubilization with Triton X-100, suggesting that membranes or vesicles are indispensable for infection, since astrovirus particles in these assays were not proteolytically activated. Preincubation of both types of vesicles with anti-Yuc8 neutralizing polyclonal antibody left a fraction of virus particles infectious, suggesting that some of these viruses (10-20%) were protected from the neutralization by the antibodies, possibly by being inside the vesicles (Fig. 5A, B and C). Accordingly, pretreatment of the vesicle fractions with detergent, allowed complete antibody neutralization of the virus particles (Fig. 5A and B), supporting the hypothesis that vesicles in these fractions are important to allow viral infection and to shield viral particles from neutralizing antibodies.

**Figure 5.**
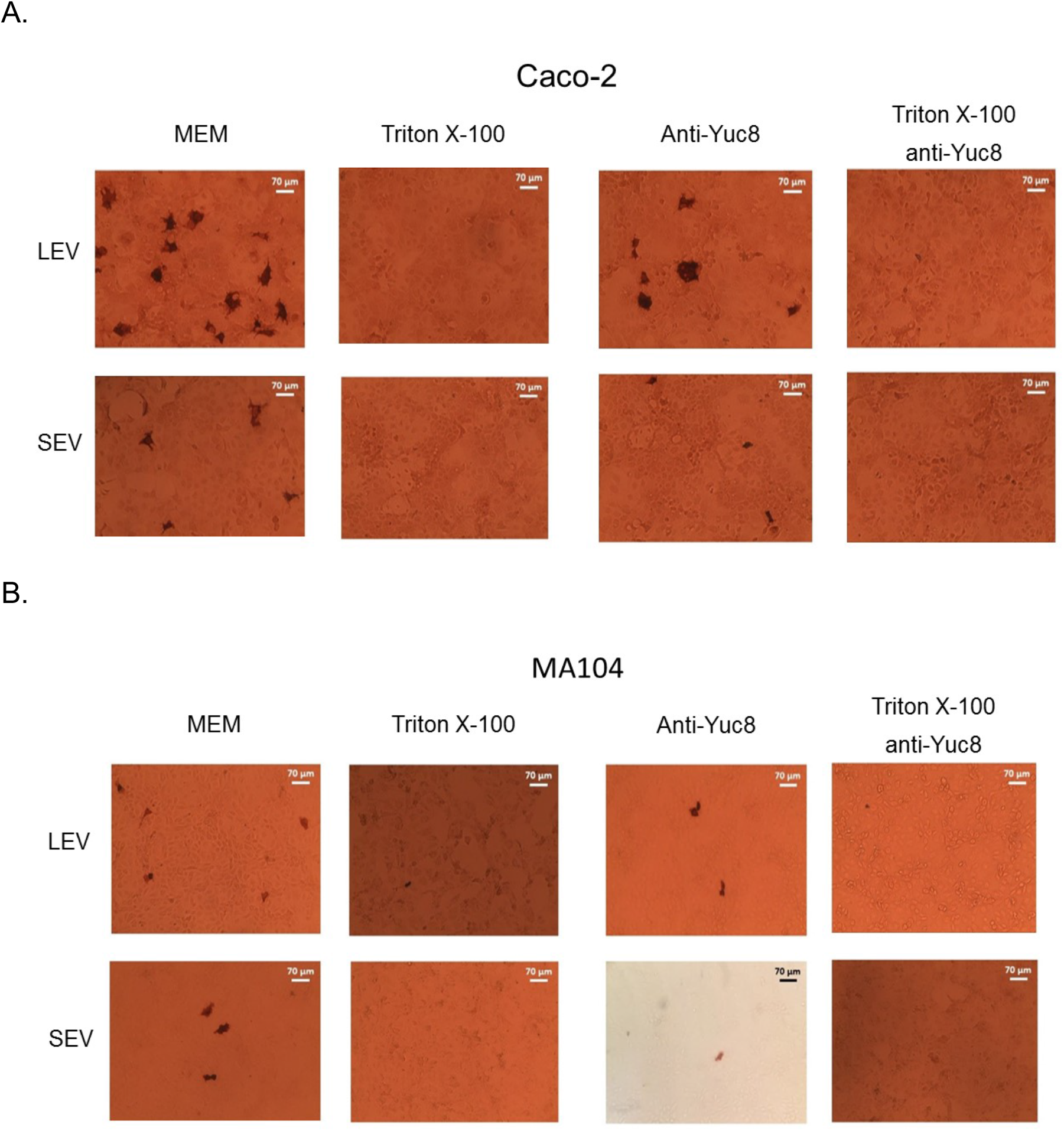
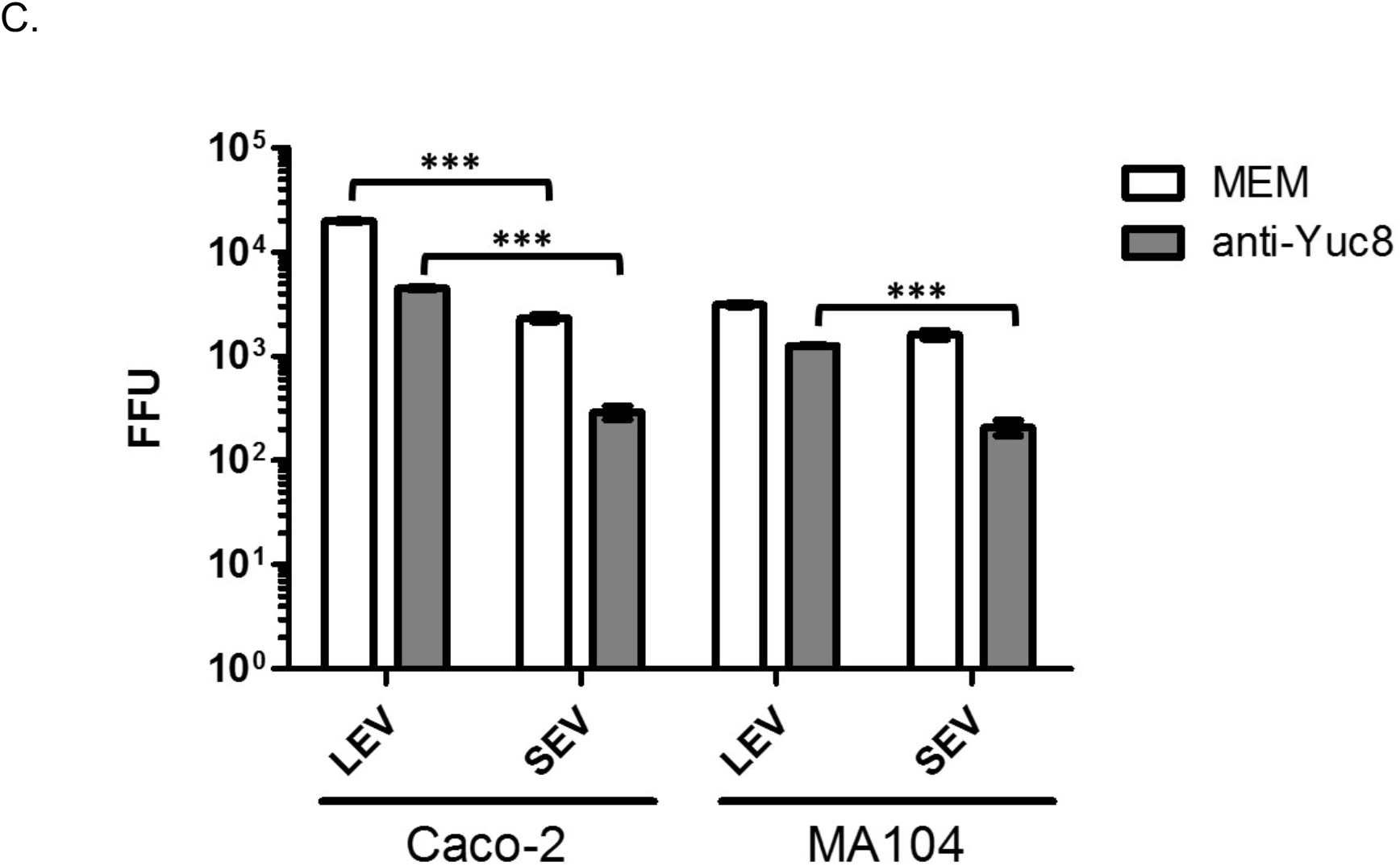
Vesicle associated astrovirus is infective in Caco-2 and MA104 cells. Large extracellular vesicles (LEV) and small extracellular vesicles (SEV) were purified by differential centrifugation and MagCapture™ exosome isolation kit PS from supernatants of Caco-2 cells infected with astrovirus (strain Yuc8). Samples were treated with medium (MEM); 0.1% Triton X-100; anti-Yuc8 (1:1500); or 0.1% Triton X-100 followed by anti-Yuc8 (1:1500) before addition to (A) Caco-2 or (B) MA104 cells. Treated samples were let to adsorb for two hours, after which time unbound vesicles were washed and infection was left to proceed for 18 hours. Infected cells were detected by immuno-peroxidase staining (darker cells). Cells were observed in a Nikon Diaphot 300 microscope with 10X magnification and they are representative of three independent experiments done in three wells each. (C) Focus forming units (FFU) of each sample were counted in 3 wells of three independent experiments. Bars represent the viral focus forming units (FFU) in each sample ± standard error of the mean. Samples where 0 FFU were observed (Triton X-100 and Triton X-100 plus anti-Yuc8 treatments) are not graphically represented. Statistical analysis was done with two-way ANOVA, p value *<0.05; ** p<0.01, *** p<0.001.

Protected viral particles were observed in both, LEV and SEV fractions, and they were able to infect a similar number of both Caco-2 and MA104 cells (Fig. 5C). When the infectivity was compared between Caco-2 and MA104 cells, Caco-2 cells showed more infected cells by LEV fraction than MA104 cells, while infection associated with SEV fraction was similar between both cell lines (Fig. 5C). In Caco-2 cells there were more infected cells after infection with vesicles from LEV fraction, as compared to SEV fraction, while in MA104 cells infectivity of these two fractions was similar (Fig. 5C).

### Association of non-activated astrovirus Yuc8 with purified EVs enhances viral infectivity

To further evaluate the possibility that the association of the virus with membranous structures promotes virus infectivity without the need of trypsin treatment, non-activated purified Yuc8 particles were incubated for 1 h at 37 °C with LEV and SEV fractions obtained from supernatants of mock-infected Caco-2 cells. After virus-EV incubation, the virus-vesicle mixture was subjected to the same treatments described in the previous experiment: neutralization with polyclonal anti-Yuc8 antibody, membrane solubilization by incubation with 0.1% Triton X-100, or detergent treatment followed by neutralization. Untreated virus-vesicle samples, in MEM, were used as control. After treatment, the samples were added to Caco-2 and MA104 monolayers and infection was left to proceed as described. The nonactivated astrovirus particles that were incubated with both types of purified vesicles (LEV and SEV fractions) acquired the capacity to infect both Caco-2 and MA104 cells (Fig. 6A and B). Detergent treatment of the samples before addition to the cells abolished the infectivity in both cell lines, again confirming the contribution of membrane vesicles to viral infectivity of particles non-activated by trypsin, possibly by direct interaction between virus and vesicles (Fig. 6A and B). Pre-incubation with neutralizing antibodies abolished infectivity, suggesting that all viral particles were accessible to the antibodies. Of note, no infection was detected when either cell line was incubated with the same amount of non-activated virus, in the absence of vesicles, unless the virus was activated by treatment with trypsin (Fig. 6C). As expected, the combined treatment of detergent and neutralizing antibodies also abolished the infection (Fig. 6A and B). These results suggest that free viral particles could associate with vesicles, and this interaction facilitates their cell entry and infectivity, even if the virus is not activated. When the infectivity of the vesicle-associated non-activated astrovirus particles was compared in Caco-2 and MA104 cells, both LEV and SEV fractions showed similar capacity to promote infection in both cell lines (Fig. 6C). Of interest, non-activated astrovirus particles incubated with purified EV, infected MA104 cells more efficiently (>200%) than the same amount of free virus activated with trypsin (Fig. 6C), while in Caco-2 cells (astrovirus fully permissive cell line), the vesicle-associated particles had a 17% average infectivity of the free, trypsin activated virus (Fig. 6C).

**Figure 6.**
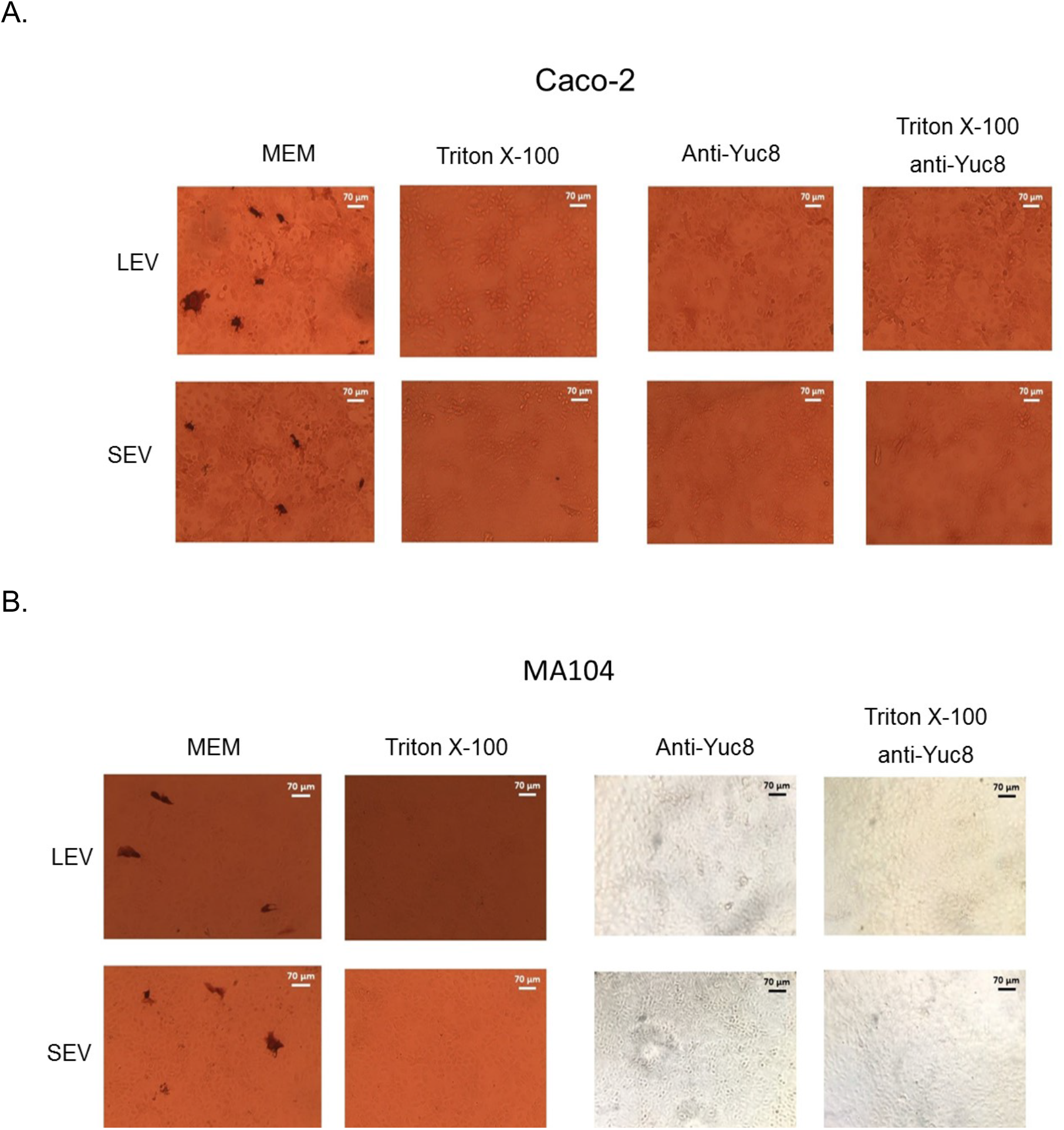
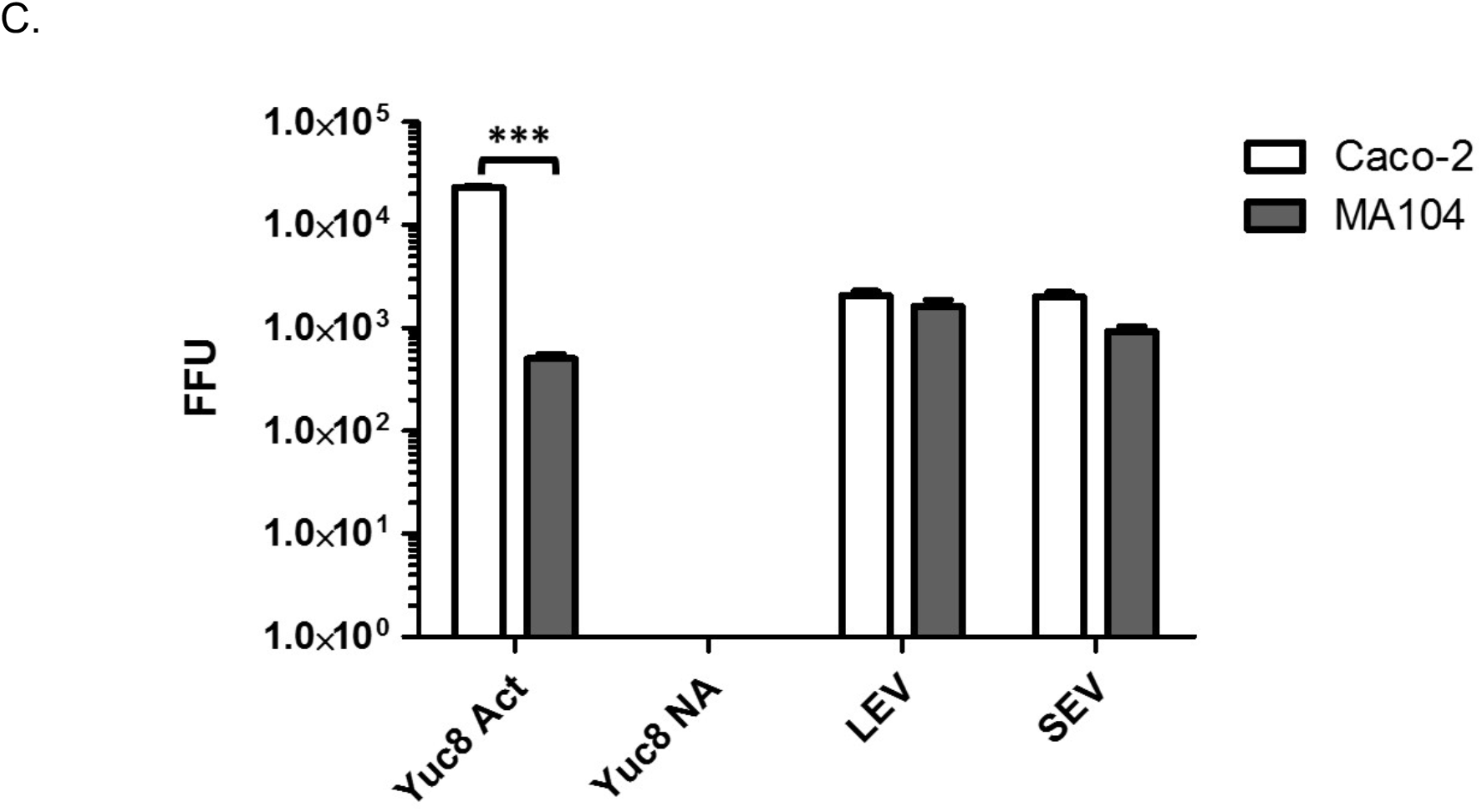
The interaction of non-activated astrovirus Yuc8 with vesicles enhances their infectivity. Large extracellular vesicles (LEV) and small extracellular vesicles (SEV) were purified by differential centrifugation and MagCapture™ exosome isolation kit PS from supernatants of non-infected Caco-2 cells. Purified vesicles were incubated with purified non-activated astrovirus Yuc8 particles for 1 hour at 37°C. Vesicle-virus mixtures were treated in 4 different conditions: medium (MEM); 0.1% Triton X-100; anti-Yuc8 (1:1500); and 0.1% Triton X-100 followed by anti-Yuc8 (1:1500). Treated fractions were then let to adsorb in (A) Caco-2 or (B) MA104 cells for two hours, after which time the unbound vesicles and viral particles were washed out. After 18 hpi infected cells were detected by immuno-peroxidase staining (darker cells). Images magnified with a 10X lens were observed in a Nikon Diaphot 300 microscope and they are representative of three independent experiments done in triplicate. (C) Infected cells observed in experimetns described in A and B were counted in three wells of three independent experiments and compared with the same amount of trypsin activated (Yuc8 Act) and non-activated (Yuc8 NA) astrovirus probed without pre-incubation with EV in Caco-2 and MA104 cells. Graphics shown the amount of infected cell in each sample expressed as focus forming units (FFU). Bars represent the mean FFU ± standard error of the mean. Statistical analysis was done with twoway ANOVA, p value *<0.05; ** p<0.01, *** p<0.001.

## Discussion

Astrovirus cell release has been reported to be a non-lytic process promoted by caspase processing of the viral capsid-precursor protein VP90 to VP70 (10, 11). It is a gradual process in which the majority of the new particles (about 90% of the total progeny) remain inside infected cells, and only 10% are released to the extracellular media (10). In this study, we found that a portion of the new progeny was present in the cell supernatant as early as 12 hpi and the amount of released virus increased with time. Interestingly, starting at 15 hpi, a significant portion of the released virus particles were not susceptible to trypsin treatment, and required to be solubilized from their association with membranous structures by detergent treatment, to become accessible to the protease. The amount of virus protected from trypsin increased with time. This observation suggests that there could be more potentially infectious virus particles in the extracellular media than originally thought (10), most probably explained by the association of viruses with EVs.

Using differential centrifugation, we purified several fractions of EV from the media of astrovirus Yuc8 infected Caco-2 cells, and two of these fractions were characterized in more detail: the LEV fraction, obtained after cellular debris depletion and by centrifugation for 1 h at 20,000 g; and the SEV fraction, purified after PEG 6000 based precipitation and centrifugation at 10,000 g (40, 42). Both fractions contained markers of extracellular vesicles, CD63 and Alix. Since CD63 is an specific marker of exosomes (50, 51), and ALIX has been reported to be involved in both microvesicle and exosome biogenesis (45), it is possible that both of these vesicles, which overlap in size, are present in both fractions. Analysis of the concentration and size of the vesicles in LEV and SEV fractions by nanoparticle tracking showed partial overlap in size, however the LEV fractions showed a significative increase in the vesicle number when infected versus mock-infected conditions were compared. A similar increase in EV secretion after infection was also reported with other viruses, like HIV-1 (52, 53), HSV-1 (33, 54), rotavirus (27) and tick-borne Langat virus (55).

Different viruses have been found to be able to interact directly with EV. Among these, hepatitis A virus (HAV) (48), HCV (29), HSV-1 (33), and JC polyomavirus (34). Analysis of the LEV fraction, purified from astrovirus-infected cells, showed electrodense astrovirus-like particles associated with vesicles of about 200 nm, resembling the appearance of extracellular vesicles (56). The membranous structures observed by TEM seem to associate with more than one viral particle. This observation opens up the possibility that during astrovirus infection, EV could participate both as virion carriers, protecting the virions, as well as a form of concentrating viral particles, forming the so called collective infectious units (CIU), capable to gather together several infectious particles. Similar observations have been made recently for rotavirus and norovirus, where several viral particles were reported to be associated with vesicles (27, 57).

It has been described previously that the association with vesicles could protect some viruses from neutralization by antibodies, for example hepatitis A, B and C viruses (29, 48, 58), and HSV-1 (33), among others. In this work we observed that a portion of the astrovirus particles present in LEV and SEV fractions remained infectious even after incubation with an anti-Yuc8 neutralizing antibody, suggesting that a portion of the isolated astrovirus particles were inaccessible to the neutralizing antibodies. The presence of vesicles was crucial for the infectivity of these nonactivated viral particles, since the solubilization of membranes with detergent abrogated all infectivity. To confirm that vesicles are important in non-activated astrovirus infectivity, LEV and SEV vesicles were purified from non-infected Caco-2 cells, and then incubated with non-trypsin activated purified astrovirus particles. Our results showed that the purified virus, was able to interact with these vesicles, and acquired the capacity to enter and infect the cells without protease activation. It is not clear if these interactions between astrovirus particle and EVs interaction are specific or not, but the virus particles in this mix acquired the capacity to infect even low susceptibility cells like MA104. The infectivity was abolished by solubilization of the vesicles with detergent, or by incubation with neutralizing antibodies, suggesting that the interaction between viral particles and EV somehow facilitates the interaction between the virus and the cell surface.

Since extracellular vesicles could facilitate the internalization of the virus apparently through a viral receptor independent pathway, the viral particles associated with vesicles could be internalized by a mechanism triggered by vesicles themselves (59). The incubation of non-activated purified astrovirus with LEV or SEV fractions leads to similar level of infection in both Caco-2 and MA104 cells, while non-activated purified astrovirus particles were not able to infect these cell lines. These results suggest that the astrovirus proteolytic processing by trypsin (activation), is important for virus-cell adhesion and/or entry, but probably not for the decapsidation process. The ability of astrovirus particles associated with vesicles to infect not only susceptible Caco-2 cells, but also the poorly susceptible MA104 cell line (Fig 6C), suggest that the vesicle-associated virus particles could bypass certain blocks in astrovirus tropism, probably the specific virus-receptor interaction, potentially increasing their pathogenicity. This observation also suggest that extracellular vesicles could help astrovirus to disseminate outside the gastrointestinal tract like it has been reported before for HAstV serotype 4 and the novel astroviruses strains MLB and VA (2), possibly by allowing astroviruses to avoid the immune response and cellular barriers until they get into permissive cells far away from their common environment (gastrointestinal tract).

Our observations suggest the possibility that EV could be acting as platforms to create collective infectious units (60, 61), rendering virus particles insensitive to neutralization with antibodies and promoting their internalization in a non-receptor dependent manner. The mechanisms by which EV promote viral internalization in new cells remain unclear, as well as the contribution of EV to the whole astrovirus infectivity.

## Acknowledgments

This work was supported by grants 254608 and 302965 from CONACyT (Mexico). We thank to Dra. Guadalupe Zavala for her assistance in transmission electron microscopy. Carlos Baez-N was recipient of scholarship from CONACyT (Mexico), No. 464741 for MS degree, and had support from the postgraduate studies program (PAEP) of the National Autonomous University of Mexico (UNAM).

## References

1. Meliopoulos V, Schultz-Cherry S. 2012. Astrovirus Pathogenesis., p 65–75. In Schultz-Cherry S (ed), Astrovirus Research. Springer, New Yorl, NY.

2. Vu DL, Bosch A, Pinto RM, Guix S. 2017. Epidemiology of Classic and Novel Human Astrovirus: Gastroenteritis and Beyond. Viruses 9.

3. Vu DL, Cordey S, Brito F, Kaiser L. 2016. Novel human astroviruses: Novel human diseases? J Clin Virol 82:56–63.

4. Bosch A, Pinto RM, Guix S. 2014. Human astroviruses. Clin Microbiol Rev 27:1048–74.

5. E. M, Arias CF. 2013. Astroviruses, p 1347–1401. In Knipe DM (ed), Fields Virology, vol 2. Lippincot Williams & Wilkins, Philadelpia.

6. Aguilar-Hernandez N, Meyer L, Lopez S, DuBois RM, Arias CF. 2020. Protein Disulfide Isomerase A4 Is Involved in Genome Uncoating during Human Astrovirus Cell Entry. Viruses 13.

7. Arias CF, DuBois RM. 2017. The Astrovirus Capsid: A Review. Viruses 9.

8. Brinker JP, Blacklow NR, Herrmann JE. 2000. Human astrovirus isolation and propagation in multiple cell lines. Arch Virol 145:1847–56.

9. Mendez E, Arias CF. 2013. Astroviruses, p 1347–1401. In Knipe DM (ed), Fields Virology. Lippincot Williams and Wilkins, Philadephia.

10. Banos-Lara Mdel R, Mendez E. 2010. Role of individual caspases induced by astrovirus on the processing of its structural protein and its release from the cell through a non-lytic mechanism. Virology 401:322–32.

11. Mendez E, Salas-Ocampo E, Arias CF. 2004. Caspases mediate processing of the capsid precursor and cell release of human astroviruses. J Virol 78:8601–8.

12. Aguilar-Hernandez N, Lopez S, Arias CF. 2018. Minimal capsid composition of infectious human astrovirus. Virology 521:58–61.

13. Mendez E, Fernandez-Luna T, Lopez S, Mendez-Toss M, Arias CF. 2002. Proteolytic processing of a serotype 8 human astrovirus ORF2 polyprotein. J Virol 76:7996–8002.

14. Mendez E, Aguirre-Crespo G, Zavala G, Arias CF. 2007. Association of the astrovirus structural protein VP90 with membranes plays a role in virus morphogenesis. J Virol 81:10649–58.

15. Altan-Bonnet N, Chen YH. 2015. Intercellular Transmission of Viral Populations with Vesicles. J Virol 89:12242–4.

16. Barteneva NS, Maltsev N, Vorobjev IA. 2013. Microvesicles and intercellular communication in the context of parasitism. Front Cell Infect Microbiol 3:49.

17. Schorey JS, Cheng Y, Singh PP, Smith VL. 2015. Exosomes and other extracellular vesicles in host-pathogen interactions. EMBO Rep 16:24–43.

18. Raposo G, Stoorvogel W. 2013. Extracellular vesicles: exosomes, microvesicles, and friends. J Cell Biol 200:373–83.

19. Minciacchi VR, Freeman MR, Di Vizio D. 2015. Extracellular vesicles in cancer: exosomes, microvesicles and the emerging role of large oncosomes. Semin Cell Dev Biol 40:41–51.

20. Thery C, Witwer KW, Aikawa E, Alcaraz MJ, Anderson JD, Andriantsitohaina R, Antoniou A, Arab T, Archer F, Atkin-Smith GK, Ayre DC, Bach JM, Bachurski D, Baharvand H, Balaj L, Baldacchino S, Bauer NN, Baxter AA, Bebawy M, Beckham C, Bedina Zavec A, Benmoussa A, Berardi AC, Bergese P, Bielska E, Blenkiron C, Bobis-Wozowicz S, Boilard E, Boireau W, Bongiovanni A, Borras FE, Bosch S, Boulanger CM, Breakefield X, Breglio AM, Brennan MA, Brigstock DR, Brisson A, Broekman ML, Bromberg JF, Bryl-Gorecka P, Buch S, Buck AH, Burger D, Busatto S, Buschmann D, Bussolati B, Buzas EI, Byrd JB, Camussi G, et al. 2018. Minimal information for studies of extracellular vesicles 2018 (MISEV2018): a position statement of the International Society for Extracellular Vesicles and update of the MISEV2014 guidelines. J Extracell Vesicles 7:1535750.

21. Thery C, Ostrowski M, Segura E. 2009. Membrane vesicles as conveyors of immune responses. Nat Rev Immunol 9:581–93.

22. Casorla-Perez LA, Lopez T, Lopez S, Arias CF. 2018. The Ubiquitin-Proteasome System Is Necessary for Efficient Replication of Human Astrovirus. J Virol 92.

23. Karpe YA, Meng XJ. 2012. Hepatitis E virus replication requires an active ubiquitin-proteasome system. J Virol 86:5948–52.

24. Kourembanas S. 2015. Exosomes: vehicles of intercellular signaling, biomarkers, and vectors of cell therapy. Annu Rev Physiol 77:13–27.

25. Lai FW, Lichty BD, Bowdish DM. 2015. Microvesicles: ubiquitous contributors to infection and immunity. J Leukoc Biol 97:237–45.

26. Altan-Bonnet N. 2016. Extracellular vesicles are the Trojan horses of viral infection. Curr Opin Microbiol 32:77–81.

27. Isa P, Perez-Delgado A, Quevedo IR, Lopez S, Arias CF. 2020. Rotaviruses Associate with Distinct Types of Extracellular Vesicles. Viruses 12.

28. Meckes DG, Jr. 2015. Exosomal communication goes viral. J Virol 89:5200–3.

29. Liu Z, Zhang X, Yu Q, He JJ. 2014. Exosome-associated hepatitis C virus in cell cultures and patient plasma. Biochem Biophys Res Commun 455:218–22.

30. Nagashima S, Jirintai S, Takahashi M, Kobayashi T, Tanggis, Nishizawa T, Kouki T, Yashiro T, Okamoto H. 2014. Hepatitis E virus egress depends on the exosomal pathway, with secretory exosomes derived from multivesicular bodies. J Gen Virol 95:2166–2175.

31. Nagashima S, Takahashi M, Kobayashi T, Tanggis, Nishizawa T, Nishiyama T, Primadharsini PP, Okamoto H. 2017. Characterization of the Quasi-Enveloped Hepatitis E Virus Particles Released by the Cellular Exosomal Pathway. J Virol 91.

32. Tamai K, Shiina M, Tanaka N, Nakano T, Yamamoto A, Kondo Y, Kakazu E, Inoue J, Fukushima K, Sano K, Ueno Y, Shimosegawa T, Sugamura K. 2012. Regulation of hepatitis C virus secretion by the Hrs-dependent exosomal pathway. Virology 422:377–85.

33. Bello-Morales R, Praena B, de la Nuez C, Rejas MT, Guerra M, Galan-Ganga M, Izquierdo M, Calvo V, Krummenacher C, Lopez-Guerrero JA. 2018. Role of Microvesicles in the Spread of Herpes Simplex Virus 1 in Oligodendrocytic Cells. J Virol 92.

34. Morris-Love J, Gee GV, O’Hara BA, Assetta B, Atkinson AL, Dugan AS, Haley SA, Atwood WJ. 2019. JC Polyomavirus Uses Extracellular Vesicles To Infect Target Cells. mBio 10.

35. Mendez-Toss M, Romero-Guido P, Munguia ME, Mendez E, Arias CF. 2000. Molecular analysis of a serotype 8 human astrovirus genome. J Gen Virol 81:2891–2897.

36. Oceguera A, Peralta AV, Martinez-Delgado G, Arias CF, Lopez S. 2018. Rotavirus RNAs sponge host cell RNA binding proteins and interfere with their subcellular localization. Virology 525:96–105.

37. Mendez E, Salas-Ocampo MP, Munguia ME, Arias CF. 2003. Protein products of the open reading frames encoding nonstructural proteins of human astrovirus serotype 8. J Virol 77:11378–84.

38. Sanchez-Fauquier A, Carrascosa AL, Carrascosa JL, Otero A, Glass RI, Lopez JA, San Martin C, Melero JA. 1994. Characterization of a human astrovirus serotype 2 structural protein (VP26) that contains an epitope involved in virus neutralization. Virology 201:312–20.

39. Guerrero CA, Zarate S, Corkidi G, Lopez S, Arias CF. 2000. Biochemical characterization of rotavirus receptors in MA104 cells. J Virol 74:9362–71.

40. Rider MA, Hurwitz SN, Meckes DG, Jr. 2016. ExtraPEG: A Polyethylene Glycol-Based Method for Enrichment of Extracellular Vesicles. Sci Rep 6:23978.

41. Thery C, Amigorena S, Raposo G, Clayton A. 2006. Isolation and characterization of exosomes from cell culture supernatants and biological fluids. Curr Protoc Cell Biol Chapter 3:Unit 3 22.

42. Livshits MA, Khomyakova E, Evtushenko EG, Lazarev VN, Kulemin NA, Semina SE, Generozov EV, Govorun VM. 2015. Isolation of exosomes by differential centrifugation: Theoretical analysis of a commonly used protocol. Sci Rep 5:17319.

43. Quevedo IR, O’sson ALJ, Clark RJ, Veinot JGC, Tufenkji N. 2014. Interpreting deposition behavior of polydisperse surface-modified nanoparticles using QCM-D and sand-packed columns. Environmental Engineering Science, 31:326–337.

44. Hassellow M, Kaegi R. 2009. Analysis and characterization of manufactured nanoparticles in aquatic environments., p 211–266. In Lead JR, Smith E (ed), Environmental and human health impacts of nanotechnology. Wiley, Chichester, West Sussex.

45. van Niel G, D’Angelo G, Raposo G. 2018. Shedding light on the cell biology of extracellular vesicles. Nat Rev Mol Cell Biol 19:213–228.

46. Mack M, Kleinschmidt A, Bruhl H, Klier C, Nelson PJ, Cihak J, Plachy J, Stangassinger M, Erfle V, Schlondorff D. 2000. Transfer of the chemokine receptor CCR5 between cells by membrane-derived microparticles: a mechanism for cellular human immunodeficiency virus 1 infection. Nat Med 6:769–75.

47. Rozmyslowicz T, Majka M, Kijowski J, Murphy SL, Conover DO, Poncz M, Ratajczak J, Gaulton GN, Ratajczak MZ. 2003. Platelet- and megakaryocyte-derived microparticles transfer CXCR4 receptor to CXCR4-null cells and make them susceptible to infection by X4-HIV. AIDS 17:33–42.

48. Feng Z, Hensley L, McKnight KL, Hu F, Madden V, Ping L, Jeong SH, Walker C, Lanford RE, Lemon SM. 2013. A pathogenic picornavirus acquires an envelope by hijacking cellular membranes. Nature 496:367–71.

49. Aguilar-Hernandez N. 2018. Caracterizacion de las interacciones tempranas de astrovirus de humano con celulas permisivas y resistentes a la infeccion. master. Universidad Nacional Autonoma de Mexico.

50. Kalra H, Simpson RJ, Ji H, Aikawa E, Altevogt P, Askenase P, Bond VC, Borras FE, Breakefield X, Budnik V, Buzas E, Camussi G, Clayton A, Cocucci E, Falcon-Perez JM, Gabrielsson S, Gho YS, Gupta D, Harsha HC, Hendrix A, Hill AF, Inal JM, Jenster G, Kramer-Albers EM, Lim SK, Llorente A, Lotvall J, Marcilla A, Mincheva-Nilsson L, Nazarenko I, Nieuwland R, Nolte-’t Hoen EN, Pandey A, Patel T, Piper MG, Pluchino S, Prasad TS, Rajendran L, Raposo G, Record M, Reid GE, Sanchez-Madrid F, Schiffelers RM, Siljander P, Stensballe A, Stoorvogel W, Taylor D, Thery C, Valadi H, van Balkom BW, et al. 2012. Vesiclepedia: a compendium for extracellular vesicles with continuous community annotation. PLoS Biol 10:e1001450.

51. Kowal J, Arras G, Colombo M, Jouve M, Morath JP, Primdal-Bengtson B, Dingli F, Loew D, Tkach M, Thery C. 2016. Proteomic comparison defines novel markers to characterize heterogeneous populations of extracellular vesicle subtypes. Proc Natl Acad Sci U S A 113:E968–77.

52. Arenaccio C, Chiozzini C, Columba-Cabezas S, Manfredi F, Affabris E, Baur A, Federico M. 2014. Exosomes from human immunodeficiency virus type 1 (HIV-1)-infected cells license quiescent CD4+ T lymphocytes to replicate HIV-1 through a Nef- and ADAM17-dependent mechanism. J Virol 88:11529–39.

53. Kulkarni R, Prasad A. 2017. Exosomes Derived from HIV-1 Infected DCs Mediate Viral transInfection via Fibronectin and Galectin-3. Sci Rep 7:14787.

54. Deschamps T, Kalamvoki M. 2018. Extracellular Vesicles Released by Herpes Simplex Virus 1-Infected Cells Block Virus Replication in Recipient Cells in a STING-Dependent Manner. J Virol 92.

55. Zhou W, Woodson M, Neupane B, Bai F, Sherman MB, Choi KH, Neelakanta G, Sultana H. 2018. Exosomes serve as novel modes of tick-borne flavivirus transmission from arthropod to human cells and facilitates dissemination of viral RNA and proteins to the vertebrate neuronal cells. PLoS Pathog 14:e1006764.

56. Rikkert LG, Nieuwland R, Terstappen L, Coumans FAW. 2019. Quality of extracellular vesicle images by transmission electron microscopy is operator and protocol dependent. J Extracell Vesicles 8:1555419.

57. Santiana M, Ghosh S, Ho BA, Rajasekaran V, Du WL, Mutsafi Y, De Jesus-Diaz DA, Sosnovtsev SV, Levenson EA, Parra GI, Takvorian PM, Cali A, Bleck C, Vlasova AN, Saif LJ, Patton JT, Lopalco P, Corcelli A, Green KY, Altan-Bonnet N. 2018. Vesicle-Cloaked Virus Clusters Are Optimal Units for Inter-organismal Viral Transmission. Cell Host Microbe 24:208–220 e8.

58. Yang Y, Han Q, Hou Z, Zhang C, Tian Z, Zhang J. 2017. Exosomes mediate hepatitis B virus (HBV) transmission and NK-cell dysfunction. Cell Mol Immunol 14:465–475.

59. McKelvey KJ, Powell KL, Ashton AW, Morris JM, McCracken SA. 2015. Exosomes: Mechanisms of Uptake. J Circ Biomark 4:7.

60. Sanjuan R. 2017. Collective Infectious Units in Viruses. Trends Microbiol 25:402–412.

61. Sanjuan R. 2018. Collective properties of viral infectivity. Curr Opin Virol 33:1–6.

